# Coordinated postnatal maturation of striatal cholinergic interneurons and dopamine release dynamics in mice

**DOI:** 10.1101/2020.04.02.022152

**Authors:** Avery McGuirt, Ori Lieberman, Michael Post, Irena Pigulevskiy, David Sulzer

## Abstract

Dynamic changes in motor abilities and motivated behaviors occur during the juvenile and adolescent periods. The striatum is a subcortical nucleus critical for action selection, motor learning and reward processing. Its tonically active cholinergic interneuron (ChI) is an integral regulator of the synaptic activity of other striatal neurons, as well as afferent axonal projections of midbrain dopamine neurons. Thalamic and dopaminergic inputs initiate pauses in ChI firing following salient sensory cues that are extended for several hundred milliseconds by intrinsic regenerative currents. Here, we characterize the electrophysiological and morphological features of ChIs during mouse postnatal development. We demonstrate that ChI spontaneous activity increases with age while the duration of the pause in firing induced by depolarizing inputs decreases during postnatal development. Maturation of ChI activity is driven by two distinct physiological changes: decreased amplitude of the afterhypolarization between P14 and P18 and and increased Ih conductance between the late postnatal period and adulthood. Finally, we uncover postnatal changes in dopamine release properties that are mediated by cholinergic signalling. At P10, striatal dopamine release is diminished compared to the adult, but our data show efficient summation of dopamine relase evoked by multiple grouped stimuli that subsides by P28. Blockade of nictonic acetylcholine receptors enhances release summation in mice older than P28 but has little effect at P10. These data demonstrate a physiological maturation of ChI activity and indicate a reciprocal interaction between the postnatal maturation of striatal ChI and dopamine neurotransmission.

**Significance Statement:** Motor skills and motivated behavior regimes develop rapidly during the postnatal period. The functional development of the striatal cholinergic interneuron (ChI), which contributes to these behaviors in adulthood, remains unexplored. In this study, we tracked the ontogeny of spontaneous ChI activity and cellular morphology, as well as the developmental trajectory of ion conductances characteristic to this population. We further report a developmental link between ChI activity and dopamine release, revealing a change in the frequency-dependence of dopamine release during the early postnatal period that is mediated by cholinergic signaling. This study provides evidence that striatal microcircuits are dynamic during the postnatal period and that they undergo coordinated maturation.

## Introduction

The early postnatal period is a time of increasing sensory perception and the development of complex motor behaviors (Altman and Sudarshan, 1975; Shaywitz et al., 1979; Westerga and Gramsbergen, 1990). In rodents, locomotor activity increases dramatically during the second postnatal week (Altman and Sudarshan, 1975; Shaywitz et al., 1979; Westerga and Gramsbergen, 1990), together with increased exploration and the acquisition of motivated behaviors that integrate internal and external states (Hall et al., 1977).

The striatum is the main input nucleus of the basal ganglia and contributes to action selection, motor learning and motivated behaviors in the adult (Gerfen and Surmeier, 2011). The cholinergic interneuron (ChI) comprises only about 1-5% of striatal neurons (Kemp and Powell, 1971a, 1971b), but due to its widespread axonal arborization and synaptic connections with other striatal neurons, acts as a critical node in striatal synaptic computation (DiFiglia et al., 1976; DiFiglia, 1987; Kawaguchi, 1993; Goldberg and Reynolds, 2011; Goldberg et al., 2012). ChIs form axo-axonic synapses with dopaminergic axons and regulate dopamine (DA) release during motivated behaviors (Le Novère et al., 1996; Azam et al., 2002; Zoli et al., 2002; Exley and Cragg, 2008; Sulzer et al., 2016; Mohebi et al., 2019). ChIs are tonically pace-making neurons with spontaneous firing frequencies between 2-10 Hz (Wilson et al., 1990; Bennett and Wilson, 1999). *In vivo*, ChIs respond to rewarding or aversive salient stimuli with pauses in firing that can last for several hundred milliseconds (Aosaki et al., 1994b). It is unknown if ChI tonic activity or the mechanisms that drive pauses in activity mature postnatally.

In the adult, spontaneous ChI activity is driven by intrinsic ion conductances that occur in the absence of synaptic activity (Bennett et al., 2000). I_h_, the current mediated by hyperpolarization-actived cyclic nucleotide-gated (HCN) channels, depolarizes ChIs to −60 mV, where HCN channels inactivate. A persistent sodium current then drives the cell to its action potential threshold where Ca_V_2 calcium channels open (Bennett et al., 2000). After the cell fires, calcium-activated potassium channels, S_K_ and B_K_ repolarize the cell and control the magnitude of a change in voltage known as the “medium afterhyperpolarization” (mAHP) (Goldberg and Wilson, 2005).

In the adult, a pause in ChI activity following salient cues is initiated by excitatory thalamic inputs (Matsumoto et al., 2001) and is dependent on DA signaling (Aosaki et al., 1994a; Reynolds et al., 2004; Zhang et al., 2018). The decrease in firing rate persists beyond the initial excitatory input and is driven by a combination of I_h_ and a barium-sensitive potassium current that may be mediated by delayed-rectifier K_V_7 potassium channels (Wilson, 2005; Zhang et al., 2018). The intrinsic component of this pause is known as the “slow afterhyperpolarization” (sAHP) and can be experimentally evoked by injecting depolarizing current.

To address the postnatal maturation of ChI activity, we performed cell-attached and whole-cell recordings of ChIs in the dorsal striatum in acute brain slices from mice over a range of ages. We found that the spontaneous activity of ChIs increases linearly from postnatal day 10 (P10) into adulthood. Two distinct transitions in ChI physiology drive the changes in firing rate: the mAHP decreases dramatically between P14 and P18, followed by an increase in the putative HCN current between P28 and adulthood. In addition to the maturation of spontaneous activity, the sAHP decreases in length from P10 to adulthood. Finally, using fast-scan cyclic voltammetry (FSCV), we show that immature DA release properties at P10 arise from the absence of striatal cholinergic tone. These data provide a foundation for further studies of the role of ChIs in the postnatal acquisition of complex motor tasks and motivated behaviors.

## Materials and Methods

### Animals

C57Bl6J breeder pairs were obtained from Jackson Laboratories (Bar Harbor, ME). Transgenic Dat-Ires-Cre Ai38 mice were generated as described (Lieberman et al., 2017). Mice were housed in same-sex groups of 2-4 on a 12-hour light/dark cycle with water and food available ad libitum. Breeding pairs were checked daily for pregnancy and new litters. Mice were used for experiments on the specified postnatal day (± 1) in all experiments. All experimental procedures were approved by the Columbia University Institutional Animal Care and Use Committee and followed NIH guidelines. Data combine male and female mice, and no differences were observed between sexes.

### Electrophysiology

Acute brain slices were generated as described previously (Lieberman et al., 2018; Lieberman et al., 2020). Mice underwent rapid cervical dislocation. The brain was placed in ice-cold cutting buffer (in mM): 10 NaCl, 2.5 KCl,25 NaHCO_3_, 0.5 CaCl_2_, 7 MgCl_2_, 1.25 NaH_2_PO_4_, 180 sucrose, 10 glucose bubbled with 95% O_2_/5% CO_2_ to pH 7.4. Coronal slices (250 μm) were generated using a Leica vibratome and placed in artificial cerebrospinal fluid (ACSF) and allowed to rest at 34°C for 30 minutes. The recipe for ACSF was (in mM): 125 NaCl, 2.5 KCl, 25 NaHCO_3_, 2 CaCl_2_, 1 MgCl_2_, 1.25 NaH_2_PO_4_ and 10 glucose bubbled with 95% O_2_/5% CO_2_ to pH 7.4. Slices were then maintained at room temperature for a maximum of 5 hours for recordings.

At the time of recording, slices were transferred to the recording chamber and superfused with ACSF maintained at 34°C. ChIs were identified based on the large soma size under IR/DIC optics using a 40X water immersion objective. Liquid junction potential was not corrected. Data were acquired using an Axon Instruments Axopatch 200, digitized using a Digidata 1440A at 10 kHz, and filtered at 5 kHz.

Cell-attached and whole cell recordings were accomplished using glass pipettes (2-6 MΩ) filled with internal solution (in mM): 115 potassium gluconate, 20 KCl, 20 HEPES, 1 MgCl_2_, 2 MgATP, 0.2 NaGTP adjusted to pH 7.25 with KOH, osmolarity 285 mOsm. Spontaneous firing frequency in cell-attached mode was recorded for 3 minutes. Subsequently, a giga-ohm seal was achieved and the cell membrane was ruptured. Spontaneous action potential firing was recorded in current clamp, followed by measurement of the IV curve. Finally, inward currents were recorded in voltage clamp mode.

### Dendritic reconstructions

Dendritic reconstructions were obtained and analyzed essentially as described (Lieberman et al., 2018). Briefly, neurobiotin (1mg/mL, Vector Laboratories) was added to the internal solution and allowed to diffuse into the cell for 15 minutes after the whole cell configuration was established. The slice was then fixed in 4% PFA in 0.1M phosphate buffer (PB), pH 7.4 overnight. Slices were stained with Alexafluor488-conjugated streptavidin (1:200, ThemoFisher) in 0.6% TritonX-100 in TBS. Finally, slices were mounted and cover slipped. Slides were imaged on a Leica SP5 confocal microscope in system optimized Z-stacks using a 20X objective. Dendritic trees were traced using the simple neurite tracer plugin in ImageJ. Traced neurites were collapsed into a max projection and analyzed using the Sholl Analysis plugin.

### Cyclic voltammetry

Electrochemical recordings of evoked DA release by fast-scan cyclic voltammetry (FSCV) were collected as detailed previously (Lieberman et al., 2018). Carbon fiber working electrodes were made by aspirating a single carbon fiber (5 μm diameter) into a glass capillary (1.2mm borosilicate, A-M Systems), and pulling to a long taper with a micropipette puller (Sutter; P-97). Fibers were cut to an exposed length of ~100 μm, and silver leads (0.015”; A-M Systems) were permanently affixed inside the pipette by coating with colloidal silver paint before insertion. Striatal slices were prepared as for electrophysiology experiments (see above). During recordings, slices were kept under constant superfusion of oxygenated ACSF (2 mL/min, 34°C). *A* carbon fiber working electrode was placed in the dorsolateral striatum approximately 50 μm into the slice. A triangular voltage wave (−450 to+800 mV at 294 mV/ms versus Ag/AgCl) was applied across the working electrode every 100 ms and current was monitored with an Axopatch 200B amplifier (Axon Instruments) using a 5 kHz low-pass Bessel Filter setting and 25 kHz sampling rate. Signals were digitized using an ITC-18 board (Instrutech) and recorded with IGOR Pro 6.37 software (WaveMetrics), using in-house acquisition procedures. Slices were stimulated with a sharpened bipolar concentric electrode (400μm max outer diameter; Pt/Ir; WPI), placed ~150 μm from the recording electrode, using an Iso-Flex stimulus isolator (AMPI) triggered by a Master-9 pulse generator (AMPI). A single stimulus pulse (100 μs × 200 μA) was applied every 2 min until stable release was achieved, after which three consecutive peaks were averaged to define single pulse release magnitude. A train stimulus was then applied (100 Hz x 5 pulses). For nAChR antagonism, slices were then superfused with ACSF containing DHßE (1 μM). Slices were again stimulated with single pulses every two minutes until stable release (generally 10-15 minutes) was achieved. Finally, slices were again stimulated with the 100 Hz train. For p10 slices that showed no obvious DHßE-evoked change in release, 100 Hz stimuli were applied at least 15 minutes after the start of DHßE perfusion. In a given recording condition, train stimulation evoked no subsequent changes in single pulse DA release (data not shown). Data were processed and peaks quantified using an in-house procedure in IGOR Pro, and summary data was analyzed using Prism 7 (GraphPad). Electrodes were calibrated by quantifying background-subtracted voltammograms in standard solutions of DA in ACSF, made fresh each recording day.

### Two-photon imaging

Two-photon images were acquired on a Prairie (Middleton, WI) Ultima microscope system using PrairieView 4.3 software. Acute brain slices from DAT-ires-Cre x GCAMP3 mice were collected as described above, transferred into a chamber, and perfused with ACSF at room temperature. Samples were excited with a Coherent (Santa Clara, CA) Chameleon Ultra two-photon laser tuned to 920 nm, and images were collected through a photomultiplier tube channel with a 490-560 nm emission filter. The objective used was a 60X, 0.9 NA water immersion lens (Olympus). Time series were acquired from a 100 x 100 pixel ROI located within in the same field of view (1024 x 1024 pixels) as the electrode, at max speed (~ 0.15 sec / frame) for 480 frames, in Galvo mode with a dwell time 8 μs. Pulses (200 μA x 100 μS) were delivered to the slice and triggered by a Master 8 pulse generator (AMPI) via a concentric bipolar electrode (WPI, see above). The slice received a single pulse or 10 pulses at 100 Hz. This was repeated ~8-12 times per slice. Each ROI was quantified for mean pixel intensity in Image J (NIH) and the first and final 5 seconds were used to fit an exponential decay to each trace. The mean intensities were then normalized to the fit to derive a baseline corrected trace to correct for photobleaching, and an average of traces was calculated for each slice, serving as each N.

### Experimental Design and Statistical Analysis

Electrophysiology data were analyzed offline using Clampfit software (Molecular Devices, Sunnyvale, California). Statistical analysis was conducted in GraphPad Prism 7 (La Jolla, CA). All bar graphs show the mean+/- standard error of the mean. Data comparing two variables was analyzed with a two-way ANOVA. Post-hoc Bonferroni tests were conducted when significant differences were found with the two-Way ANOVA. Data comparing one variable among >2 groups was analyzed with one-Way ANOVA and Bonferroni post-tests and among 2 groups a two-tailed t test. Data were not formally tested for parametric distribution. Group sizes were preliminarily determined based on past work (Lieberman et al., 2018).

## Results

### The firing patterns and frequency of ChIs mature postnatally

To address how the activity of ChIs matures postnatally, we performed cell attached recordings of visually identified ChIs at a range of ages across postnatal development. Cell-attached recordings were utilized to assess spontaneous activity in order to minimally disturb the intracellular milieu and preserve spontaneous activity. ChIs were visually identified by their large cell bodies under DIC optics and cellular identity was further confirmed following conversion to the whole cell configuration (see below). We recorded from a total of 100 ChIs at P10, P14, P18, P28 and adults (P110-P120). Two cells recorded from a mouse at P10 were not spontaneously active in the cell-attached configuration but showed classic ChI characteristics in the whole-cell configuration (discussed below) and were included in subsequent analyses. All spontaneously active cells were considered to fire with tonic or rhythmic firing patterns except for two cells (one recorded at P14 and one at P18) which were considered to be “bursty” (Bennett and Wilson, 1999).

Spontaneous firing frequencies significantly increased across postnatal development (Figure 1*A,B*). ChIs exhibit variation in the regularity of their firing and ChIs classically exhibit an inverse correlation between spontaneous firing frequency and the coefficient of variation of their firing in the acute brain slice (Bennett and Wilson, 1999). Consistently, the coefficient of variation of ChIs decreased during postnatal development (Figure 1*C*). When the firing frequency and coefficient of variation is plotted for each individual recorded cell, a clear inverse correlation is observed between these parameters (Figure 1*D*). We thus conclude that the spontaneous activity of ChIs matures postnatally.

**Figure 1.**
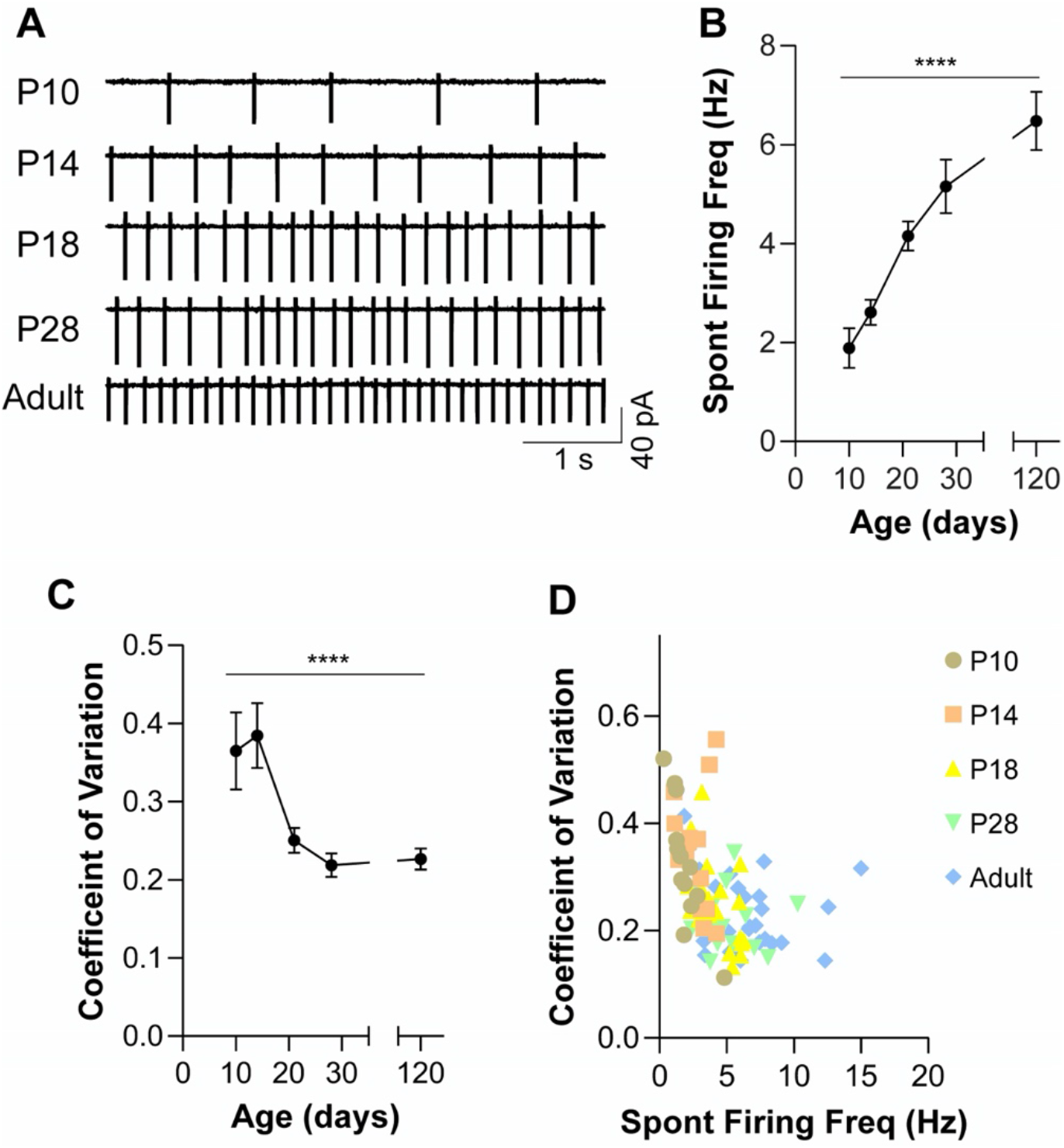
Postnatal maturation of spontaneous ChI activity. **(A)** Sample cell-attached recordings from ChIs at P10, P14, P18, P28 and adults (P110-120). **(B)** A significant increase in ChI spontaneous firing frequency from P10 to adulthood. P10 n = 18 cells (4 mice), P14 n = 16(2), P18 n = 23(4), P28 n = 15(3), Adult n=28(6). N is the same in **(C-D). (C)** Age significantly affects the coefficient of variation of ChI spontaneous activity. **(D)** Plot of coefficient of variation and spontaneous firing frequency for each recorded cell. Cells are colored by age. R^2^ = 0.205; p<0.0001. **** p<0.0001. Data analyzed in **(B)** and **(C)** with one-way ANOVA. Mean +/- sem is shown.

### ChI dendritic arborization is mature by P10

Neuronal firing patterns can be influenced by the complexity of their dendritic arbor (Mainen and Sejnowski, 1996). Following cell-attached recordings, a whole-cell configuration was established and internal solution containing neurobiotin (1 mg/mL) was allowed to diffuse into the cell. Sections were post-fixed, stained with fluorescently-labelled streptavidin and the filled cells were reconstructed using confocal microscopy. A subset of 61 cells from the 100 recorded as described above were successfully reconstructed. Cumulative dendritic length was not significantly different between the ages examined (Figure 2*A*). Interestingly, Sholl analysis revealed a transient significant increase in dendritic complexity in ChIs at P14 compared to P10, returning by P18 (Figure 2*B,C*). These data suggest that the dendritic arborization of ChIs is largely mature by P10 and reveal a previously unreported stage of dendritic overgrowth and regression at P14.

**Figure 2.**
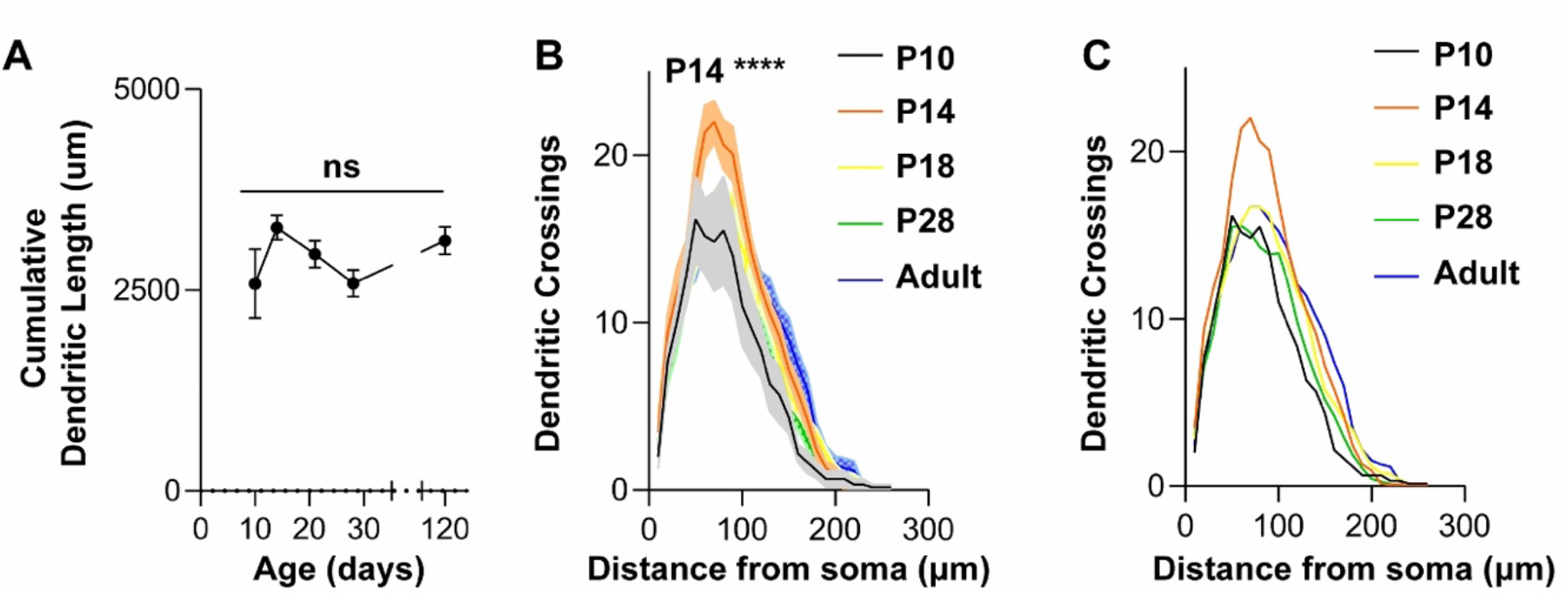
Postnatal maturation of ChI dendritic morphology. **(A)** There was no effect of age on the cumulative dendritic length of ChIs. P10: n = 6(3), P14: n = 13 (2), P18:15 (4), P28: n = 14(3), Adult: n = 13(5). **(B)** Sholl analysis reveals a significantly increased number of dendritic crossings in ChIs from P14 mice 50-90 μm from the soma compared to all other ages. Data analyzed by two-way repeated measures ANOVA followed by post-hoc Bonferroni test. Age x Distance: F(84, 1232) = 1.430, p=0.008. **(C)** Mean dendritic crossings are shown without error bars for clarity. Data are the same as in **(B)**.

### Maturation of the mAHP, but not resting potential or action potential threshold, occurs during the juvenile period

As the dendritic arborization of ChIs does not mature in parallel with spontaneous firing frequency, we examined additional properties of ChI physiology that could contribute to the increasing firing frequency observed during postnatal development.

First, we confirmed that the age-dependent increase in ChI spontaneous firing frequency also occurred in the whole-cell configuration (data not shown). The recorded cells were observed to exhibit classic features of ChIs including a sag in response to hyperpolarizing current injection and a pause in firing following depolarizing current injection (Figure 3*A*) (Kawaguchi, 1993; Bennett and Wilson, 1999). Cells without a sag or pause were excluded from the dataset.

**Figure 3.**
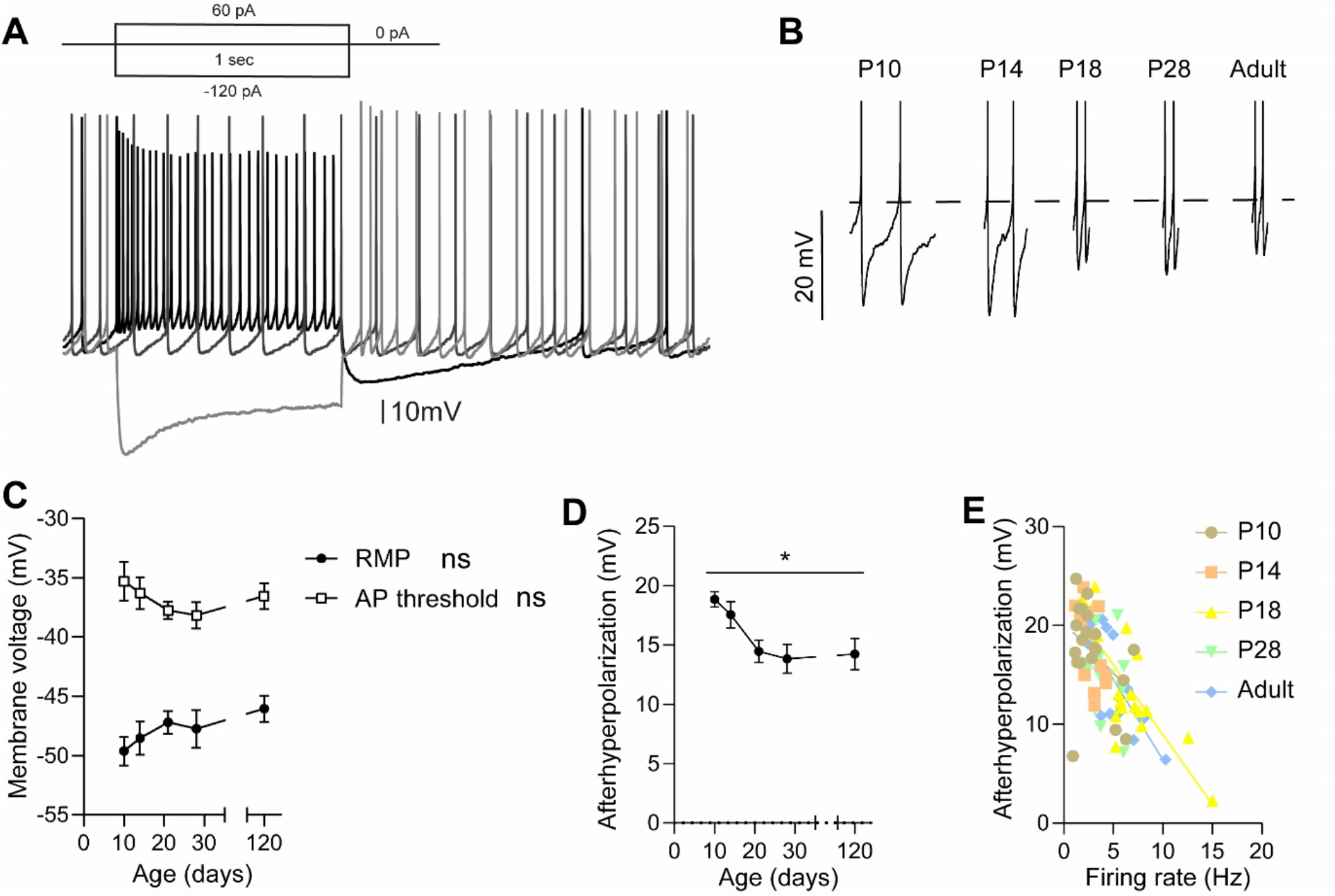
ChI afterhyperpolarization decreases between P14 and P18. **(A)** Sample current clamp trace of ChI activity in response to depolarizing and hyperpolarizing current injections. **(B)** Sample expanded traces of pairs of action potentials from ChIs recorded at the specified ages. Dashed line denotes action potential threshold. **(C)** No significant effect of age on resting membrane potential (RMP) (age: p = 0.2919) or AP threshold (age: p = 0.4404). **(D)** A significant effect of age on the afterhyperopolarization (p = 0.0102). For **(C-D)**, P10 n = 18 cells (4 mice), P14 n = 16(2), P18 n = 23(4), P28 n = 15(3), Adult n = 28(6). **(E)** Correlation between afterhyperpolarization and firing rate in ChIs from all ages. R^2^ = 0.433; p<0.0001. **(C-D)** analyzed with a one-way ANOVA.

We analyzed properties of ChI action potentials that could underlie changes in firing frequency. Sample pairs of action potentials are shown in Figure 3*B*. In these examples, the action potential threshold did not significantly differ with age (Figure 3*C*). To emphasize the salient features of these traces, the action potential amplitude is truncated. Neither action potential width nor amplitude was significantly affected by age (data not shown).

One possible explanation for an increased spontaneous firing frequency of ChIs during postnatal development is that the action potential threshold becomes more hyperpolarized, allowing the threshold to be reached more readily. Alternatively, the resting membrane potential (reported as the midpoint between two action potentials) could become more depolarized. We found, however, that age did not significantly affect either parameter (Figure 3*A,B*).

In contrast, the magnitude of the mAHP significantly decreased between P10 and adulthood (Figure 3*C*). Further, a robust negative correlation is observed between mAHP and firing rate in ChIs (Figure 3*D*), suggesting that decreasing mAHP may contribute to the developmental increase in spontaneous firing frequency. The mAHP reached the adult level by P18, however, indicating that additional changes in ChI physiology drive increases in firing frequency that occur beyond this age.

### Maturation of inward currents

Depolarization of ChIs from the hyperpolarized potential of the mAHP is driven by I_h_ (Bennett et al., 2000; Robinson and Siegelbaum, 2003). The magnitude of I_h_ can be measured in voltage clamp experiments by holding the cell at −60 mV, where HCN channels are closed, and stepping the cell to hyperpolarized potentials (Bennett et al., 2000). I_h_ inactivates following a hyperpolarizing voltage step and the current remaining at steady-state is mediated by leak channels (Figure 4*A*). Thus the magnitude of the inactivated current, or difference between maximal and steady-state inward current following hyperpolarizing voltage step, is a proxy for I_h_ (Robinson and Siegelbaum, 2003). This inactivated inward current was stable between P10 and P28 but was significantly increased in adulthood (Figure 4*B*). These data suggest that increased I_h_ may contribute to elevated spontaneous firing frequencies in adulthood.

**Figure 4.**
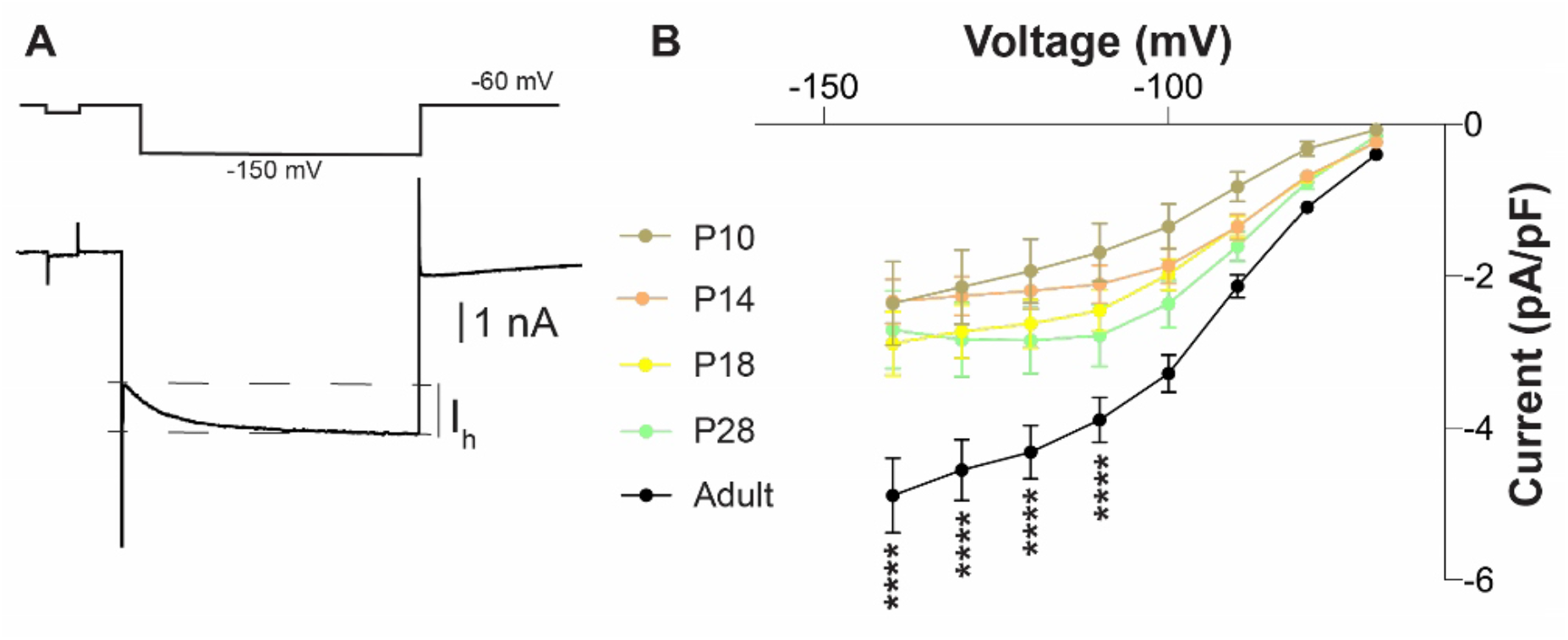
Postnatal maturation of putative I_h_ currents. **(A)** Sample voltage clamp recording demonstrating the inactivating I_h_ current. **(B)** Aggregate Ih current density shows a significant increase between P28 and adulthood but no difference at younger ages. P10 n = 18 cells (4 mice), P14 n = 16(2), P18 n = 23(4), P28 n = 15(3), Adult n = 28(6). Data analyzed by two-way repeated measure ANOVA followed by Bonferroni post-hoc test. Age x current: F(28,602)=4.580, p<0.0001.

### The sAHP is extended during the early juvenile period

A unique feature of ChI physiology is a pause in firing in response to salient sensory stimuli (Aosaki et al., 1994b). Although the pause can be initiated by excitatory thalamic inputs (Matsumoto et al., 2001) or D2 receptor activation (Aosaki et al., 1994a; Reynolds et al., 2004; Ding et al., 2010; Straub et al., 2014), intrinsic potassium and non-specific cation (I_h_) conductances define its duration (Wilson, 2005; Wilson and Goldberg, 2006; Zhang et al., 2018). These conductances can also be triggered by injecting depolarizing currents, which mimic innervation by synaptic inputs or DA (Wilson, 2005; Wilson and Goldberg, 2006; Zhang et al., 2018) and drive a pause in firing, or sAHP. A depolarizing current step was delivered to ChIs at different ages and the duration of the pause in firing after the end of the current injection was measured (Figure 5*A*). The duration of the sAHP was longer at P10 and P14 than in adulthood (Figure 5*A,B*). We conclude that the conductances that drive the sAHP are exaggerated during the early juvenile period.

**Figure 5.**
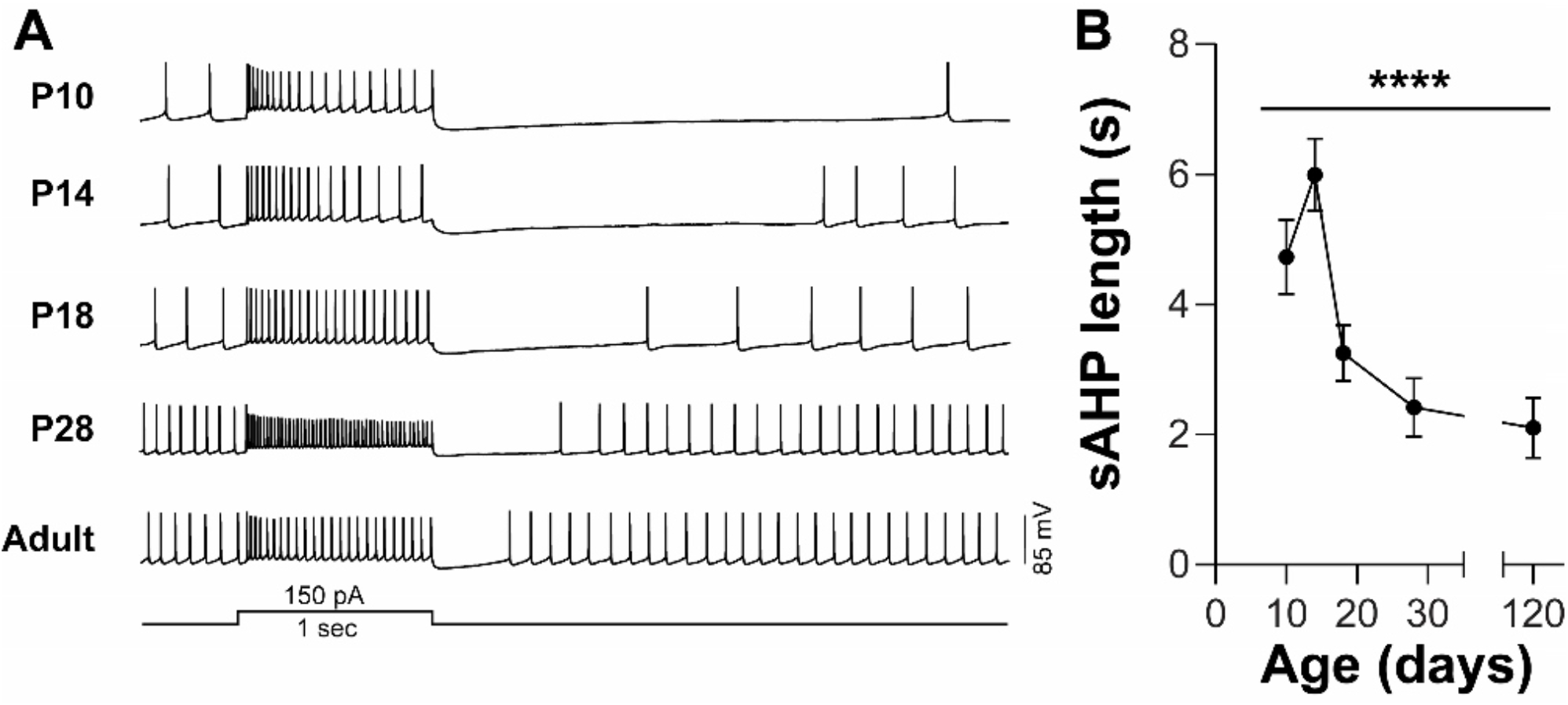
Postnatal decreases in the sAHP. **(A)** Sample current clamp recording showing the response to injection of depolarizing current. The length of the pause in firing following the end of the current step is quantified in **(B)**. **(B)** Aggregate sAHP length shows a significant effect of age. P10 n = 18 cells (4 mice), P14 n = 16(2), P18 n = 23(4), P28 n = 15(3), Adult n = 28(6). Data analyzed with one-way ANOVA.

### Summation of DA release in response to multiple grouped stimuli

The pause in ChI firing following salient sensory cues is widely considered to affect local DA release in the striatum. Acetylcholine released from striatal ChIs promotes local DA release via activation of nicotinic acetylcholine receptors (nAChR) on DA axon terminals. The interaction between DA and ACh can be probed in the acute brain slice, where it has been shown that nAChR activation on DA axons acts as a frequency-dependent filter on DA release (Zhou et al., 2001; Rice and Cragg, 2004; Zhang and Sulzer, 2004). ACh amplifies DA release in response to low-frequency stimuli, whereas prolonged stimulation or robust exogenous nAChR activation (for example, by nicotine) causes nAChR desensitization and thus decreases evoked DA release. We recently reported that the juvenile striatum shows relatively low levels of DA release in response to single electrical pulses (Lieberman et al., 2018). The changes in ChI activity over development led us to hypothesize that the low levels of ChI activity in the juvenile striatum (~P10) may lead to differences in DA release properties.

To address this, DA release was measured using fast-scan cyclic voltammetry (FSCV) in acute brain slices from mice at P10, P28, and in adulthood, following intrastriatal electrical stimulation. Consistent with our recent report, DA release following a single pulse increased significantly with age (Figure 6*A,B*). We found, however, that a train of stimuli significantly increased evoked DA release as compared to a single stimulus at P10 but not at P28 or in adulthood (Figure 6*B,C*). These results are reported as the ratio of evoked DA following 100 Hz stimuli to a single pulse (100Hz/1p; Figure 6*C*), and the ratio is significantly lower at P28 and in adulthood compared to P10. We thus conclude that DA release properties mature between P10 and P28.

**Figure 6.**
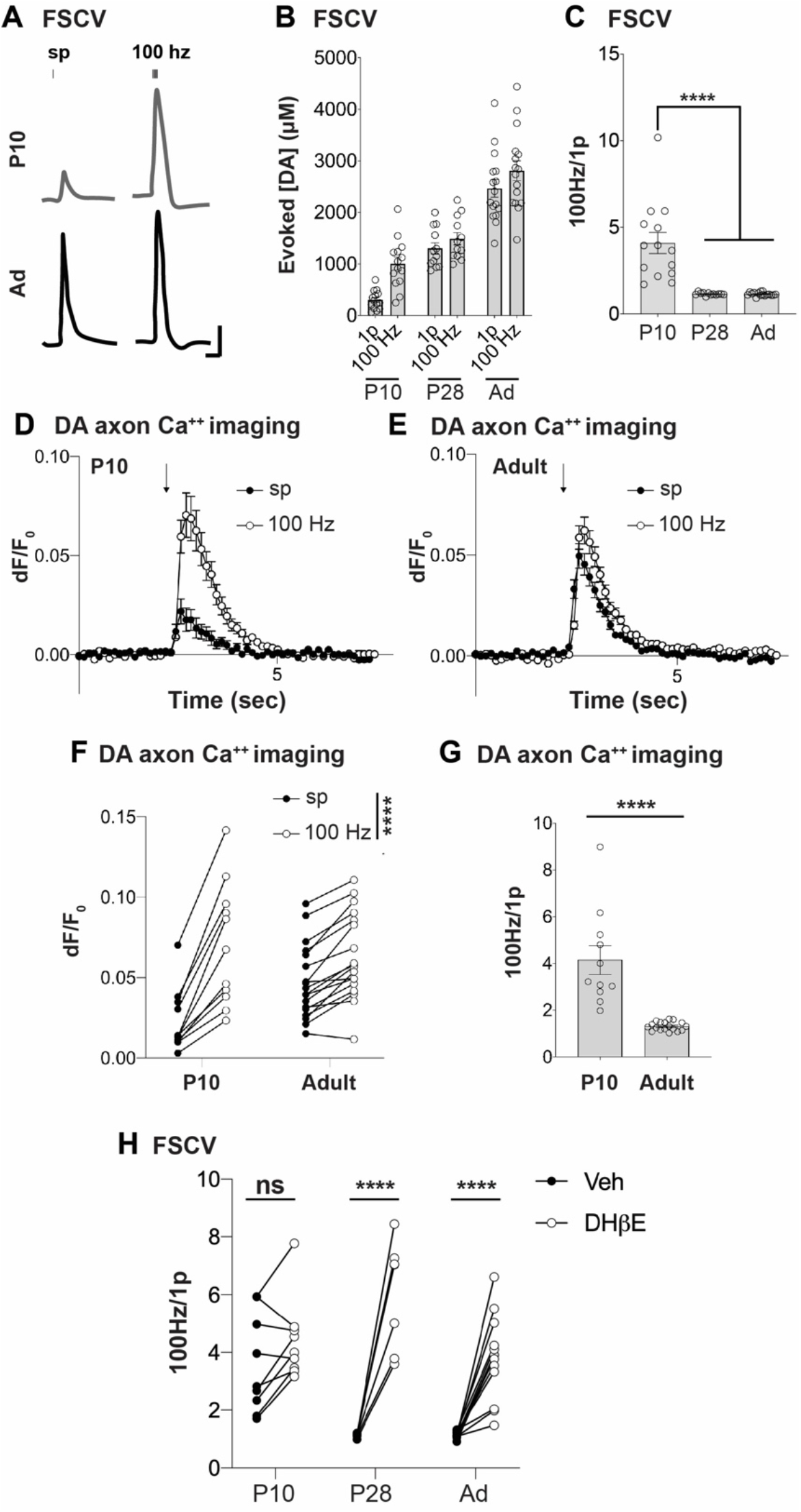
Reduced cholinergic activity at P10 leads to immature DA release properties. **(A)** Sample cyclic voltammetry recordings of evoked striatal DA release with a single pulse (sp) or 5 pulses at 100 Hz at P10 or in adult striatal slices. Scale bar: 200 nM and 200 ms. **(B)** Absolute peak concentrations of evoked DA release following a single pulse (1p) or five pulses at 100 Hz. **(C)** Absolute concentrations from **(B)** displayed as the ratio of DA evoked with 5 pulses at 100 Hz to a single pulse in striatal slices from mice aged P10, P28 or Adult. P10 n = 14 slices from 4 mice, P28 n = 12 slices from 3 mice, Adult n = 14 slices from 3 mice. Data analyzed by one-way ANOVA followed by Bonferroni post-hoc test. **(D-E)** ΔF/F of GCaMP3 in DA axons within acute slices from **(D)** P10 or **(E)** adult mice. Arrow indicates time of electrical stimulation with either a single pulse or pulses at 100 Hz. **(F)** ΔF/F from each slice after a single pulse or 100 Hz stimulation. Two-way repeated measures ANOVA: age x frequency, F_(1,27)_=40.04, p<0.0001. **(G)** Fold change in ΔF/F after 100 Hz stimulation compared to a single pulse. T_27_=5.902, p<0.0001. P10: n = 11 slices from 3 mice. Adult: n = 18 slices from 6 mice. **(H)** DHβE (1 μM) significantly increases the 100 Hz/1p ratio at P28 and adult but not at P10. Data analyzed with two-way repeated measures ANOVA followed by Bonferroni post-hoc test. Age x Drug: F(2,26) = 15.26, p<0.0001. P10 n = 9 slices from 4 mice, P28 n = 7 slices from 2 mice, Adult n = 13 slices from 3 mice.

The dynamics of neurotransmitter release are heavily dependent on calcium influx and handling. To address whether calcium entry differed in DA axons between P10 and adulthood, we generated transgenic mice expressing the calcium indicator, GCaMP3, in DA axons by crossing the DAT-Ires-Cre driver line (Bäckman et al., 2006) with the Ai38 line harboring a floxed allele of GCaMP3 in the Rosa26 locus (Zariwala et al., 2012; Lieberman et al., 2017). We confirmed specific GCaMP3 expression in DA neurons and axons at P10 and in adulthood (data not shown; (Lieberman et al., 2017)). To test whether calcium dynamics were distinct at P10 compared to adulthood, we generated acute brain slices and imaged GCaMP3 fluorescence using two-photon microscopy. A single electrical stimulation in the striatum yielded a smaller change in fluorescence at P10 compared to DA axons in the adult striatum (Figure *6D-E*).Remarkably, stimulation with electrical pulses at 100 Hz yielded an equivalent change in fluorescence at P10 and in adults (Figure *6D-E*). There was a significant interaction between age and stimulation, with DA axons at P10 having a significantly increased summation of depolarizing stimuli compared to adults (Figure *6F-G*). These data suggest that the postnatal maturation of DA release dynamics may arise from changes in calcium dynamics.

Interestingly, the observation that similar levels of DA are evoked following 100 Hz stimulation at P10 and P28 (Figure 6*B*) suggests that the lower evoked DA at P10 following a single pulse may not arise from low DA stores but rather from differences in the regulation of DA release properties particular to the immature striatal microcircuit. In the adult, the relative absence of DA release summation arises from nAChR activation on DA axons (Sulzer et al., 2016). Given the apparent efficiency of DA release summation following grouped stimuli in the juvenile striatum, we hypothesized that signaling through the nAChR is diminished in the juvenile. We thus tested the effect of a nAChR antagonist, dihydro-β-erythroidine hydrobromide (DHßE) on single-stimulus release magnitude and release summation following grouped stimuli. Evoked DA release was first measured in response to a single pulse and a train of five pulses (100 Hz), after which DHßE was superfused onto the slice and evoked DA was measured in response to the same stimulus paradigms. DHßE significantly increased the 100Hz/1p ratio at P28 and in the adult but had no effect at P10 (Figure *6H*). These data suggest that the difference in DA release properties between P10, P28, or adults arises from increasing cholinergic activity in the striatum during early postnatal maturation.

## Discussion

Here, we investigated the ontogeny of ChI firing during the postnatal development of the striatum. Using patch-clamp electrophysiology and FSCV, we have demonstrated profound changes in ChI physiology and cholinergic modulation of DA release during early postnatal development and into adulthood. Notably, significant transitions in ChI physiology occur around P14, a time when significantly more complex motor patterns are expressed (Altman and Sudarshan, 1975). The results complement previous reports showing neurochemical maturation of cholinergic signaling during this window (Coyle and Campochiaro, 1976; Coyle and Yamamura, 1976; Sawa and Stavinoha, 1987) and suggest a possible link between ChI physiology in the juvenile striatum and maturation of an animal’s behavioral repertoire.

### The development and maturation of the striatal cholinergic system

The striatum contains the highest concentration of ACh in the brain (Fibiger, 1982). In contrast to most other brain regions, however, striatal cholinergic innervation largely arises from a population of interneurons rather than from afferent innervation, for example from the basal forebrain system (Kimura et al., 1980; Henderson, 1981).

ChIs are generated between E13 and E15 (Phelps et al., 1989), prior to the appearance of striatal projection neurons or non-cholinergic striatal interneurons (E15-P2) (van der Kooy and Fishell, 1987; Song and Harlan, 1994; Liao et al., 2008). Although a major biosynthetic enzyme for ACh, choline acetyltransferase (ChAT), is present in ChIs at birth, ChAT expression levels increase linearly during the first four postnatal weeks and reach adult levels around P28 in both rat and mouse (Guyenet et al., 1975; Coyle and Yamamura, 1976; Phelps et al., 1989; Aznavour et al., 2003). Striatal ACh levels follow a similar developmental trajectory (Coyle and Campochiaro, 1976; Coyle and Yamamura, 1976; Phelps et al., 1989). The protracted postnatal maturation of ACh neurochemistry has been suggested to underlie a delayed interaction between cholinergic and dopaminergic pharmacology in the developing rodent, which reportedly reaches functional maturity around P20 (Burt et al., 1982; Fitzgerald and Hannigan, 1989). Although striatal ChIs appear in tandem with basal forebrain cholinergic neurons, their functional development lags behind the basal forebrain system (Phelps et al., 1989), suggesting that cues specific to the striatum may contribute to ChI maturation.

Here, we extend the analysis of the ontogeny of the striatal cholinergic system by electrophysiological analysis of ChIs during mouse postnatal development. We find that the maturation of ChI firing frequencies mirrors the time course of tissue ACh levels and expression of ChAT, suggesting that ACh itself may provide a feedback mechanism to drive maturation of ChI physiology during postnatal development. We also find that the time course of the maturation of ChI activity is similar to that of DA release, suggesting a possible reciprocal connection between DA signaling and ChI maturation.

### Biophysical mechanisms of the maturation of ChI firing

The frequency of tonic ChI firing is determined by the coordinated activity of intrinsic conductances (Wilson et al., 1990; Bennett et al., 2000; Goldberg and Wilson, 2005; Goldberg and Reynolds, 2011). What drives the maturation of ChI spontaneous firing described here?

At P10 and P14, the amplitude of the mAHP is significantly increased relative to ages above P18 (Figure *3*). In the adult, the mAHP is determined by the activity of S_K_ and B_K_ channels (Goldberg and Wilson, 2005). S_K_ and B_K_ currents are mediated not only by channel levels and function, but also in response to changes in calcium entry during action potential firing that initiates S_K_ and B_K_ channel opening. Future efforts will determine how the activities of these channels are altered during postnatal maturation.

A second mechanism must drive further increases in spontaneous ChI firing beyond P18, when the mAHP amplitude reaches adult levels (Figures *1* and *3*). We found that the magnitude of I_h_, which depolarizes ChIs from the mAHP toward the action potential threshold (Kawaguchi, 1993; Bennett et al., 2000), increased from P28 to adulthood (Figure *4*). Thus, the maturational increase in ChI tonic firing appears to be due to the sequential decrease in mAHP followed by an increase in I_h_.

We also report a maturational increase in the ChI pause with age. Although the ChI pause is initiated by thalamic inputs and requires DA signaling, regenerative intrinsic properties determine its duration (Aosaki et al., 1994a; Matsumoto et al., 2001; Reynolds et al., 2004; Wilson, 2005; Zhang et al., 2018). The increased pause duration over maturation is consistent with the maturation of the sAHP during postnatal development (Figure *5*). We note that for the purposes of this study, we limited analysis to the intrinsic components the ChI pause. Future work will address whether thalamic and DA inputs to ChIs mature in parallel with intrinsic mechanisms.

As the duration of the ChI pause contributes to striatal-based learning by altering DA neurotransmission and synaptic plasticity onto SPNs (Goldberg et al., 2012), differences in ChI pause dynamics might provide a mechanism for changes in learning strategy and reward sensitivity during juvenile and adolescent periods (Sturman et al., 2010; Doremus-Fitzwater et al., 2012; Sturman and Moghaddam, 2012; Spear, 2013).

### Maturation of DA release properties depends on ChIs

To address the functional implications of immature ChI activity during the juvenile period, DA release properties, which are regulated by nAChR activity in adulthood (Zhou et al., 2001; Rice and Cragg, 2004; Zhang and Sulzer, 2004; Sulzer et al., 2016), were measured using FSCV. We previously reported that evoked DA release increases during postnatal development in the striatum (Lieberman et al., 2018a). In the adult striatum, electrical stimuli elicit DA release through two mechanisms: cell-autonomous depolarization of DA axons, and via activation of nAChRs on DA axons by ACh (Zhou et al., 2001; Rice and Cragg, 2004; Zhang and Sulzer, 2004; Sulzer et al., 2016). Notably, nAChR antagonists block the majority of evoked DA after a single electrical stimulus in the adult striatum. Reduced cholinergic tone in the striatum would, thus, contribute to decreased evoked DA following a single pulse during the early juvenile period.

While acute activation of the nAChR facilitates evoked release, prolonged stimulation or pharmacological intervention cause desensitization of the nAChR and a resulting reduction in DA release capacity through this mechanism. Thus, while dopamine release in the adult is largely independent of stimulation frequency or number of grouped stimuli at baseline, nAChR desensitization elicits a capacity for summation of DA release with multiple stimuli at high frequency (Zhou et al., 2001; Rice and Cragg, 2004; Zhang and Sulzer, 2004; Sulzer et al., 2016). Here, we find that in contrast to these previous reports from adult mice, DA release is efficiently summed across multiple grouped stimuli at P10 even without pharmacological intervention. We confirmed that, in addition to DA release itself, electrically-evoked increases in calcium within DA axons also underwent postnatal maturation as DA axons at P10 had significantly more summation compared to the adult. Moreover, a nAChR antagonist did not enhance the summation of DA release at P10, but did so at P28 and in adults. We conclude that altered DA release properties in the juvenile striatum, including decreased release to a single stimulus and increased summation of DA release across grouped stimuli, are mediated by deficient signaling through the nAChR that then matures over the early postnatal period.

A limitation of this study is that we did not correlate changes in ChI firing patterns with ACh release. It is notable that tissue ACh, choline acetyltransferase, and acetylcholinesterase (AChE) each increase during postnatal development (Guyenet et al., 1975; Butcher and Hodge, 1976; Coyle and Yamamura, 1976; Murrin and Ferrer, 1984) with a time course that mirrors the maturation of spontaneous ChI firing we report. Recently developed fluorescent sensors can measure changes in extracellular ACh levels (Jing et al., 2018) and may contribute to further analysis of thse relationships. We note that a non-exclusive additional mechanism related to the regulation of DA release by ChI could be related to maturational changes in the expression of the amount, type, coupling or cell types expressing striatal nAChRs.

## Conclusion

In the adult, ChIs are widely reported to play a central role in the striatal control of action and motor learning. We report that key features of ChI electrophysiology, including spontaneous firing frequency and pauses in their activity, mature during the first four postnatal weeks, a period of acquisition of complex motor skills and enhanced sensitivity to reward. This occurs in tandem with changes in DA release properties that are mediated by cholinergic signaling, indicating that ChI and DA neurons may play reciprocal roles in the developmental regulation striatal function.

## Acknowledgments

A.F.M. was supported by NIMH (3T32NS064928-09); O.J.L. was supported by NIMH (5F30MH114390-02); MRP was supported by NIMH (5T32MH020004). This work was supported from grants by the National Institutes of Health (NIDA R01DA007418); the Simons Foundation (SFARI 514813); and the JPB Foundation. We thank Sejoon Choi for technical guidance and generous training on electrophysiology technique. We also thank Eugene Mosharov, who developed our cyclic voltammetry acquisition and analysis software and provided further technical support.

